# Hindlimb Immobilization Impairs Neuromuscular Junction Transmission in Young Rats

**DOI:** 10.1101/2025.10.15.682704

**Authors:** Nathan R. Kerr, Jose A. Viteri, Fereshteh B. Darvishi, Charles D. Brennan, Anna R. Dashtmian, Hiroshi Nishimune, Sue C. Bodine, W. David Arnold

## Abstract

**Background:** Immobilization, bed rest, or illness rapidly lead to weakness out of proportion to muscle atrophy. Although the contribution of muscle wasting to weakness is well described, the role of neuromuscular junction (NMJ) dysfunction in early disuse-related weakness is not well understood.

**Objective:** We investigated whether short-term unilateral hindlimb immobilization (HLI) in rats impairs NMJ transmission and contributes to muscle weakness out of proportion to atrophy.

**Methods:** Four-month-old male Fischer-344/Brown Norway rats underwent 10 days of unilateral HLI (n=6) or remained mobile (n=6). Neuromuscular excitability and transmission were assessed using compound muscle action potentials (CMAP), repetitive nerve stimulation (RNS), and single-fiber electromyography (SFEMG). Muscle contractility testing quantified tetanic torque, and post-mortem analysis measured muscle mass.

**Results:** Ten days of HLI reduced gastrocnemius, soleus, and plantaris muscle mass by ∼20–35%. Plantarflexion peak tetanic torque normalized to body weight declined by 23%, and torque-time integral was reduced by 36%, indicating disproportionate functional loss (muscle size versus contractile output) and supporting underlying neural impairment. CMAP amplitude decreased from 69 mV to 53.23 mV (a 22.9% reduction; p = 0.0109), indicating a loss of summated neuromuscular excitability. Furthermore, both RNS and SFEMG indicated features consistent with NMJ transmission defects. RNS revealed CMAP decrement from ∼0% pre-HLI to −9.15% at 40 Hz stimulation and −8.63% at 50 Hz post-HLI. Similarly, SFEMG confirmed marked NMJ transmission defects, with jitter increasing 103% and blocking increased from <1% to >11% of fibers.

**Conclusions:** Our findings suggest that short-term immobilization produces rapid and pronounced impairments in NMJ transmission that contribute to weakness beyond the degree of muscle atrophy. These findings identify the NMJ as an early and vulnerable site of disuse-induced dysfunction and highlight the potential for synaptic-targeted therapies to preserve muscle performance during immobilization and recovery.

## 1. Introduction

Skeletal muscle disuse, whether due to immobilization, bed rest, or critical illness, leads to rapid declines in muscle mass and function^1-3^. Within days of unloading, muscle fibers atrophy and force-generating capacity diminishes, contributing to the acceleration of physical decline, frailty, and loss of independence in older adults^4,5^. While the structural loss of muscle is well documented, it has become increasingly clear that weakness during disuse cannot be explained by atrophy alone^6,7^. Accumulating evidence suggests that neural mechanisms, including impaired excitability and neuromuscular transmission, play a key role in driving early functional decline^6,8-10^.

The neuromuscular junction (NMJ) is a critical site of communication between motor neurons and muscle fibers, ensuring reliable transmission of action potentials from nerve to muscle in order to generate sustained muscle contraction and force production. Structural and functional instability of the NMJ has been implicated in aging and neurodegenerative disease^11,12,13^, where increased NMJ jitter and blocking impair force output. Disuse-induced NMJ dysfunction, however, remains poorly understood. Our prior work in immobilized young female rats provided data suggesting the occurrence of NMJ transmission deficits based on repetitive nerve stimulation (RNS) decrement measured by electromyography (EMG)^14^. Additional animal studies have reported morphological changes at the NMJ with unloading^15,16^, yet direct electrophysiological evidence that short-term disuse impairs neuromuscular transmission is lacking. Addressing this gap is critical, as NMJ dysfunction may explain why weakness occurs disproportionately relative to muscle size loss during periods of disuse.

In the present study, we examined the effects of 10 days of unilateral hindlimb immobilization on muscle mass, function, and NMJ transmission in young male rats. We hypothesized that immobilization would induce not only measurable muscle atrophy but also impair neuromuscular excitability and transmission, thereby producing a functional deficit out of proportion to the degree of atrophy. To test this, we combined muscle contractility testing with EMG and single fiber EMG (SFEMG) to assess NMJ transmission fidelity. Our results provide direct evidence that immobilization rapidly compromises NMJ transmission, establishing synaptic deficits as a contributor to disuse-induced muscle dysfunction and weakness.

## 2. Materials and Methods

### 2.1 Animals and Experimental Design

All procedures performed herein followed NIH guidelines and received approval by the Institutional Animal Care and Use Committee of the University of Missouri. Male Fischer-344/Brown Norway rats were obtained from Charles River at 3 months of age and acclimated to our facilities for ∼3 weeks prior to the start of experiments. At ∼4 months of age, rats underwent baseline *in vivo* neuromuscular electrophysiology and muscle contractility assessments and were separated into equally balanced groups based on these results. Rats were assigned to one of two groups: fully mobile controls (CTRL, N=6) or unilateral Hindlimb Immobilization of the right hindlimb (HLI, N=6) for ten days. On day ten, casts were removed and a battery of *in vivo* electrophysiology measures were performed to investigate neuromuscular excitability, muscle contractility, and NMJ transmission as depicted in **Figure 1**. Following these measures, rats were sacrificed, muscle wet weights were measured, and tissues were collected for downstream molecular analysis.

**Figure 1:**
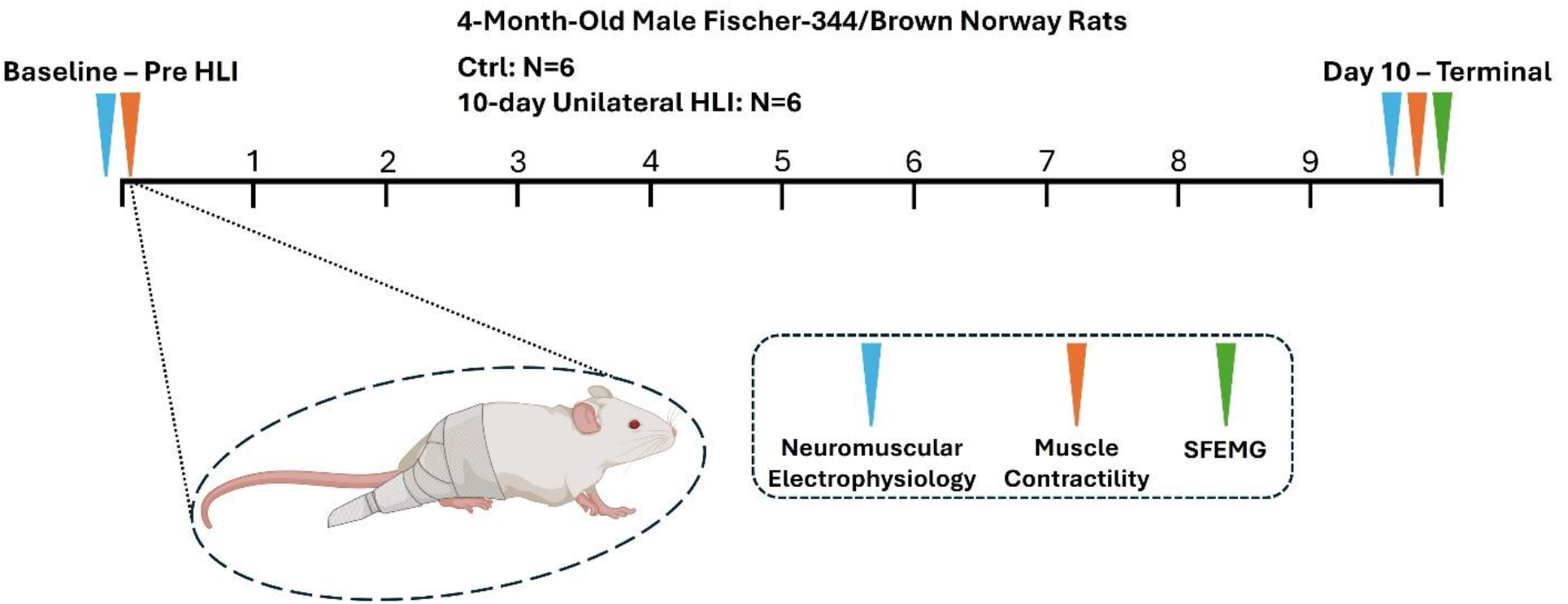
Experimental outline. 4-month-old male Fischer-344/Brown Norway rats underwent neuromuscular assessments, with an emphasis on NMJ transmission, pre- and post-unilateral hindlimb immobilization for 10 days with mobile rats serving as controls. *Figure partially created in BioRender.com*.

### 2.2 Unilateral Immobilization of the Right Hindlimb

Casts were applied after completion of baseline neuromuscular electrophysiology and muscle contractility assessments. Rats were anesthetized using isoflurane, and casting material (BSN Medical Delta-Lite Plus Casting Tape) was wrapped starting at the torso and extending down the right hindlimb with feet positioned in plantarflexion with knee at or near full extension. After limb immobilization, rats were dual housed within-group with *ad libitum* food and water access.

### 2.3 Measurement of Muscle Contractility

Immediately following neuromuscular electrophysiology assessments, muscle physiology was performed as previously described^17^. Briefly, isoflurane anesthetized rats were placed supine with the right hindlimb taped to a 90° angled foot plate attached to a force transducer. Stimulating electrodes were placed subcutaneously for supramaximal stimulation of the tibial branch of the sciatic nerve. Both twitch (a single supramaximal 0.2 ms square-wave pulse) and tetanic (a 500 ms train at 150 Hz) forces were measured to assess gastrocnemius contractile function.

### 2.4 Neuromuscular Electrophysiology Measurements of the Gastrocnemius

In vivo assessments of neuromuscular physiology were performed pre- and post-HLI using the Sierra Summit EMG system (Cadwell Sierra Summit, Kennewick, WA, USA) in a similar manner as described previously^14,17^. Briefly, rats were anesthetized using isoflurane (5% for initial sedation followed by 2% maintenance dose) and the right hindlimb was shaved, placed into full extension, and taped in place. Disc electrodes (Natus Neurology, Middleton, WI) were coated with electrode gel; the active disc electrode was placed on the gastrocnemius muscle and the reference disk electrode was placed on the metatarsal portion of the foot. A ground electrode (Cadwell) was placed on the tail. Stimulating electrodes (28-gauge monopolar needle electrodes, Teca, Oxford Instruments Medical, NY, USA) were placed subcutaneously near the sciatic nerve and stimulations were delivered while gradually increasing the stimulation intensity to achieve a maximal compound muscle action potential (CMAP) response. Supramaximal stimulation was delivered (120-150% of maximum) to record maximum CMAP which was quantified using peak-to-peak amplitude. CMAP responses during repetitive nerve stimulation (RNS) were recorded to assess neuromuscular junction (NMJ) transmission fidelity similar to our prior work in aged rats^17^. Here a train of 10 supramaximal stimuli was delivered at frequencies of 10, 20, 30, 40, and 50 Hz to evaluate the ability of the NMJ to maintain consistent CMAP amplitudes across repeated activations. The following amplifier settings were used: high pass filter: 10 kHz, low pass filter: 10 Hz, and notch filter: 60 Hz.

As an additional more direct assessment of NMJ transmission, Single Fiber Electromyography (SFEMG) was performed post-HLI using the Sierra Summit EMG system in a similar manner to our prior studies^17,18^. Due to the invasiveness of this procedure, SFEMG was only performed at the terminal timepoint. SFEMG is a clinical measure allowing for the highly sensitive assessment of NMJ transmission fidelity and its ability to generate muscle fiber action potentials^19^. Stimulating electrodes are placed subcutaneously to stimulate the sciatic nerve at a frequency of 10 Hz. Next, the SFEMG recording electrode was inserted into the right gastrocnemius muscle parallel to the muscle fibers and positioned to isolate and record single-fiber action potentials (SFAPs). Nine to ten synapses were assessed from each animal with N=3 animals per group. Jitter, or the variation in latency between action potential arrival at the nerve terminal and the generation of the corresponding SFAP. Blocking was calculated as the percentage of junctions in each animal displaying NMJ transmission failure to generate SFAPs.

### 2.5 Tissue Collection

Following final electrophysiology, SFEMG, and muscle physiology assessments under isoflurane, rats were perfused with isotonic saline, hindlimb muscles harvested and weighed, and tissues fixed or frozen in liquid nitrogen for subsequent molecular analysis.

### 2.7 Statistical Analysis

All statistical analyses were conducted using GraphPad Prism 10.2.0. For all two factor comparisons (Group: Ctrl vs HLI and Time: Pre-vs Post-), a 2-way ANOVA was used, followed by post-hoc Šidák correction analysis when significant interactions and main effects were detected. All data for these analyses met the assumption of normality as assessed by Shapiro-Wilk tests. Data sets consisting of only two comparisons (Ctrl vs HLI) were analyzed using a Student’s t-test. For data with unequal variance, a Welch’s t-test was used for these analyses instead. All data are presented with individual values as either line graphs or bar graphs (mean ± SEM) and *p* values <0.05 were considered significant.

## 3. RESULTS

### 3.1 Ten days of unilateral HLI Produces Body Weight Loss and Muscle Atrophy

4-month-old male Fischer-344/Brown Norway rats underwent unilateral HLI of the right hindlimb (N=6) for ten days with mobile rats (N=6) serving as controls. HLI resulted in a significant decrease in body weight over time (255.3 vs. 212.3 grams, p<0.0001) and relative to controls (262.8 vs. 212.3 grams, p<0.0001) while control rats gained a small amount of body weight over time (255.7 vs 262.8 grams, p=0.0306) (significant 2-way ANOVA interaction [F (1, 10) = 213.5; p<0.0001], main effect: group [F (1, 10) = 9.249; p<0.0001], and main effect: time [F (1, 10) = 110.0; p<0.0001], **Fig 2A**). Next, muscle wet weights were collected from the immobilized hindlimb and compared to controls. We noted a significant 35.25% decrease in gastrocnemius wet weight (Student’s t-test, 1201 vs 777.7 mg, t=6.23, df=10, p<0.0001, **Fig. 2B**) and a 19.74% decrease in gastrocnemius wet weight normalized to body weight (BW) (Student’s t-test, 4.575 vs 3.672 mg/g BW, t=2.877, df=10, p=0.0165, **Fig. 2C**) for HLI rats relative to controls. We noted a similar decline of 36.93% in soleus wet weight (Student’s t-test, 111.4 vs 70.27 mg, t=4.473, df=10, p=0.0012, **Fig. 2D**) and a 21.61% decline in soleus wet weight normalized to body weight (Student’s t-test, 0.4226 vs 0.3313 mg/g BW, t=2.613, df=10, p=0.0259, **Fig. 2E**). The plantaris muscle wet weight also declined by 33.6% (Student’s t-test, 258.1 vs 171.4 mg, t=4.507, df=10, p=0.0011, **Fig. 2F**) with a 17.64% decline in normalized wet weight (Student’s t-test, 0.9789 vs 0.8063 mg/g BW, t=2.821, df=10, p=0.0181, **Fig. 2G**). The extensor digitorum longus (EDL) wet weight decreased by 21.6% (Student’s t-test, 113.9 vs 89.3 mg, t=5.771, df=10, p=0.0002, **Fig. 2H**), but the normalized wet weight was unchanged (Student’s t-test, 0.4341 vs 0.4213 mg/g BW, t=0.6389, df=10, p=0.5373, **Fig. 2I**).

**Figure 2:**
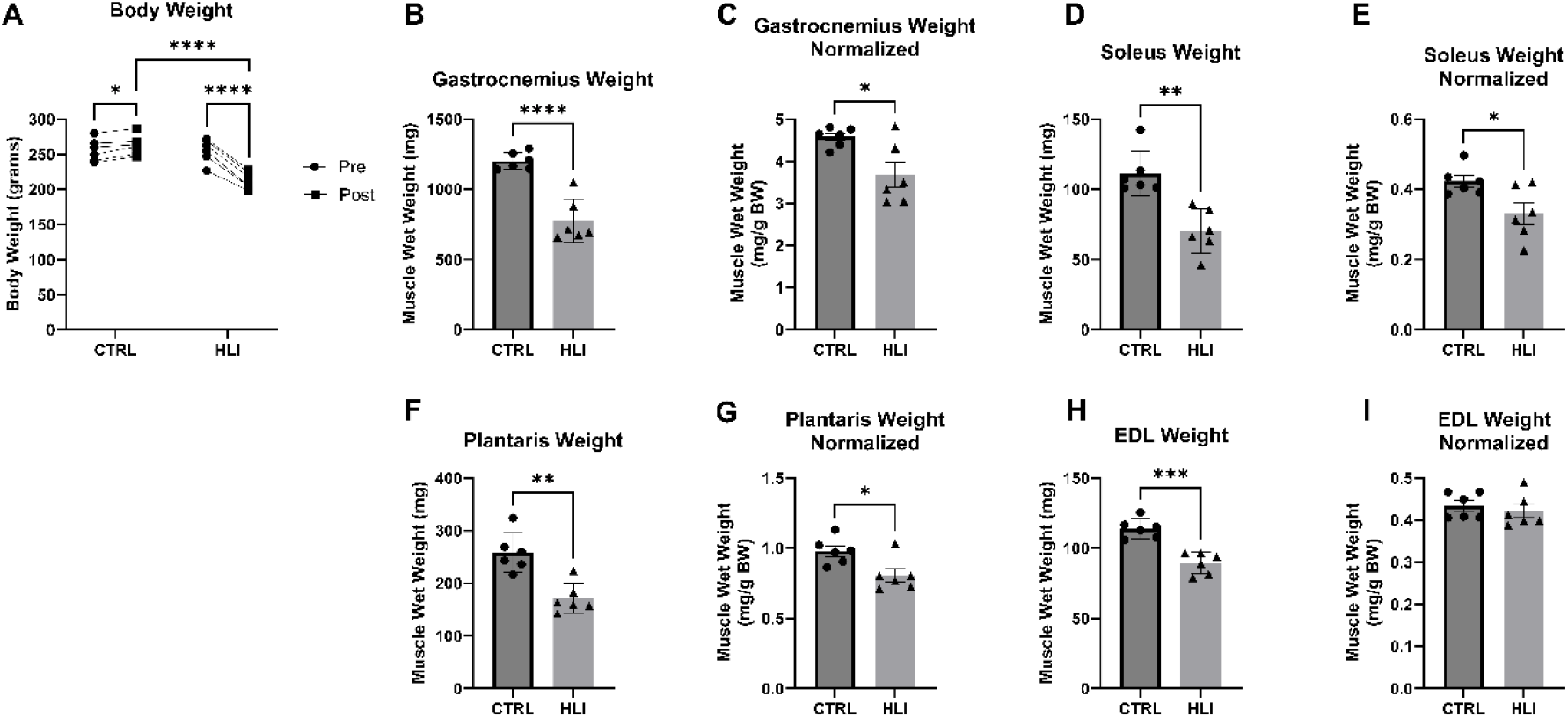
Ten days of unilateral HLI produces body weight loss and muscle atrophy. A) Body weight (BW) pre- and post-HLI. B) Gastrocnemius wet weight post-HLI and normalized to bodyweight (C). D) Soleus wet weight post-HLI and normalized to bodyweight (E). F) Plantaris wet weight post-HLI and normalized to bodyweight (G). H) EDL wet weight post-HLI and normalized to bodyweight (I). * p<0.05; ** p<0.01; *** p<0.001; ****p<0.0001.

### 3.2 Ten days of unilateral HLI Produces Loss of Muscle Contractility

Pre- and post-HLI, we assessed tetanic contractility of the gastrocnemius muscle at 150 Hz alongside mobile controls. Absolute tetanic contractility was significantly reduced post-HLI by 36.18% relative to both pre-HLI (156.7 vs 100.0 mN-m, p=0.0002) and control values (159.4 vs 100.0 mN-m, p=0.0001) (significant 2-way ANOVA interaction [F (1, 10)= 17.83, P=0.0018], main effect: group [F (1, 10) = 9.989 P=0.0101], and main effect: time [F (1, 10) = 18.80, P=0.0015], **Fig. 3A**). These differences were maintained when normalizing torque values to BW, with a significant 23.33% reduction in normalized torque values post-HLI relative to both pre-HLI (0.6135 vs 0.4704 mN-m/g BW, p=0.0037) and control values (0.6060 vs 0.4704 mN-m/g BW, p=0.0033) (significant 2-way ANOVA interaction [F (1, 10) = 6.808, P=0.0261], main effect: group [F (1, 10) = 6.513 P=0.0288], and main effect: time [F (1, 10) = 11.09, P=0.0076], **Fig. 3B**).

**Figure 3:**
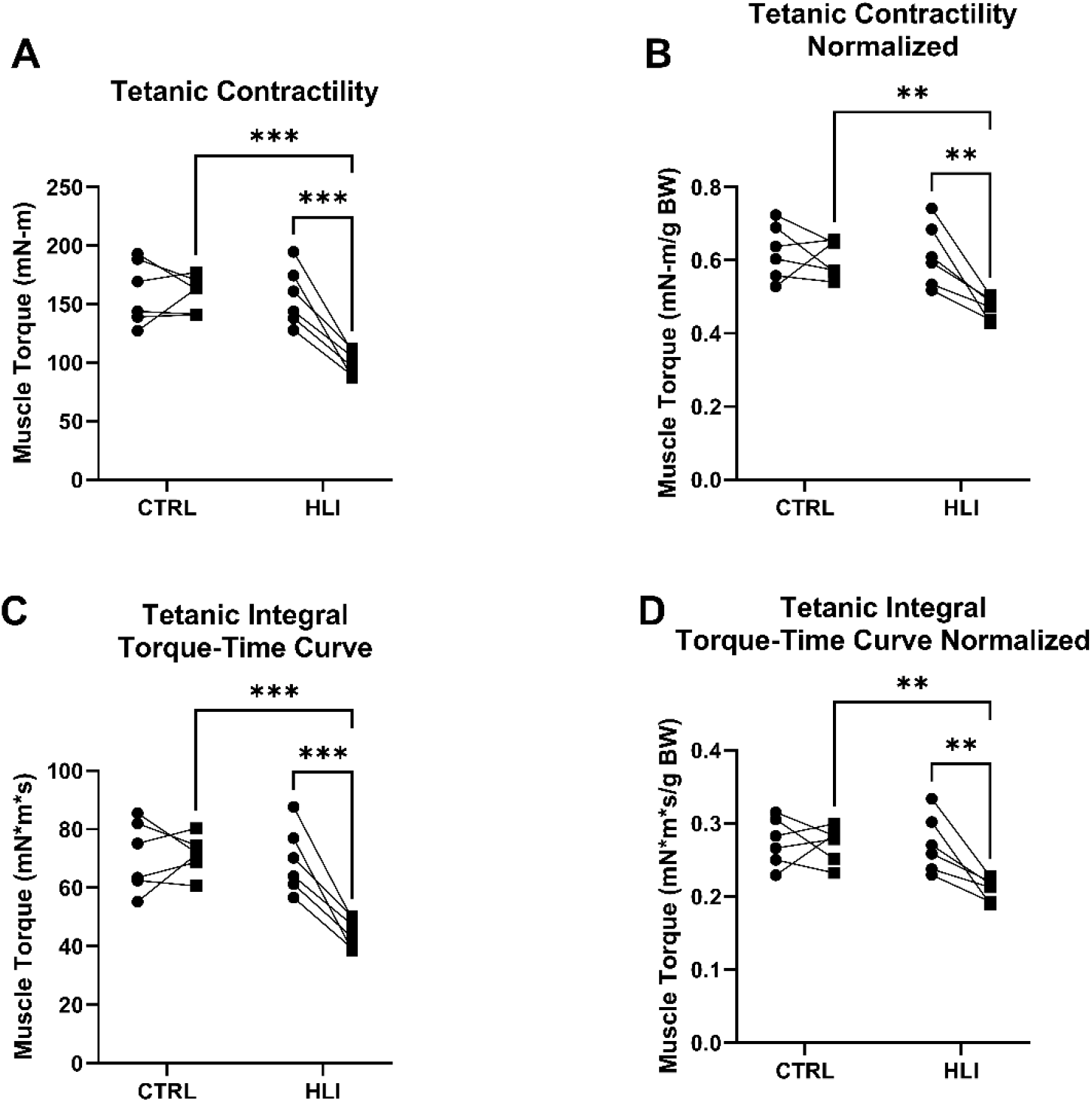
Ten days of unilateral HLI produces loss of plantarflexion muscle contractility. A) Absolute and (B) normalized gastrocnemius muscle tetanic contractility. C) Absolute and (D) normalized integral of the torque-time curve for tetanic contractility. * p<0.05; ** p<0.01; *** p<0.001.

Further, we assessed the integral of the torque-time curve to evaluate the potential contribution of NMJ transmission deficits in reduced sustained muscle contractility. Interestingly, relative to the reduction in normalized muscle torque, we found a robust, 35.71%, reduction in the integral of the torque-time curve post-HLI relative to both pre-HLI (69.43 vs 44.64 mN*m*s, p=0.0004) and control values (71.27 vs 44.64 mN*m*s, p=0.0001) (significant 2-way ANOVA interaction [F (1, 10) = 17.54, P=0.0019], main effect: group [F (1, 10) = 9.973, P=0.0102], and main effect: time [F (1, 10) = 15.76, P=0.0026], **Fig. 3C**). We noted a 22.81% reduction in the normalized integral of the torque-time curve post HLI relative to both pre-HLI (0.2719 vs 0.2099 mN*m*s/g BW, p=0.0056) and control values (0.2716 vs 0.2099 mN*m*s/g BW, p=0.0035) (significant 2-way ANOVA interaction [F (1, 10) = 6.93, P=0.0251],main effect: group [F (1, 10) = 6.195, P=0.0320], and main effect: time [F (1, 10) = 8.559, P=0.0152], **Fig. 3D**).

### 3.3 Ten days of unilateral HLI Produces Deficits in Neuromuscular Excitability and NMJ transmission

To investigate the role of NMJ transmission deficits in the observed loss of muscle contractility following HLI, we performed neuromuscular electrophysiology assessments pre- and post-HLI complemented by terminal SFEMG experiments on the gastrocnemius muscle of HLI and control rats. We began by assessing CMAP, a measure of muscle excitability, and noted a significant 22.86% CMAP amplitude decrease (69 vs 53.23 mV, p=0.0109) pre-versus post-HLI (significant 2-way ANOVA main effect: time [F (1, 10) = 8.996, P=0.0134], and trending interaction [F (1, 10) = 3.951, P=0.0749], **Fig 4A**). We then used repetitive nerve stimulation (RNS) as a measure for NMJ transmission and noted significant increases in amplitude decrement following 10 days HLI at both 40 (0.3 vs -9.15 % amplitude decrement, p=0.0019) and 50 (-1.7 vs -8.633 % amplitude decrement, p=0.0148) Hz (significant 2-way ANOVA interaction [F (4, 48) = 2.784, P=0.0369], main effect: group [F (1, 48) = 6.374, P=0.0149], and main effect: frequency [F (4, 48) = 8.417, P<0.0001], **Fig. 4B**).

**Figure 4:**
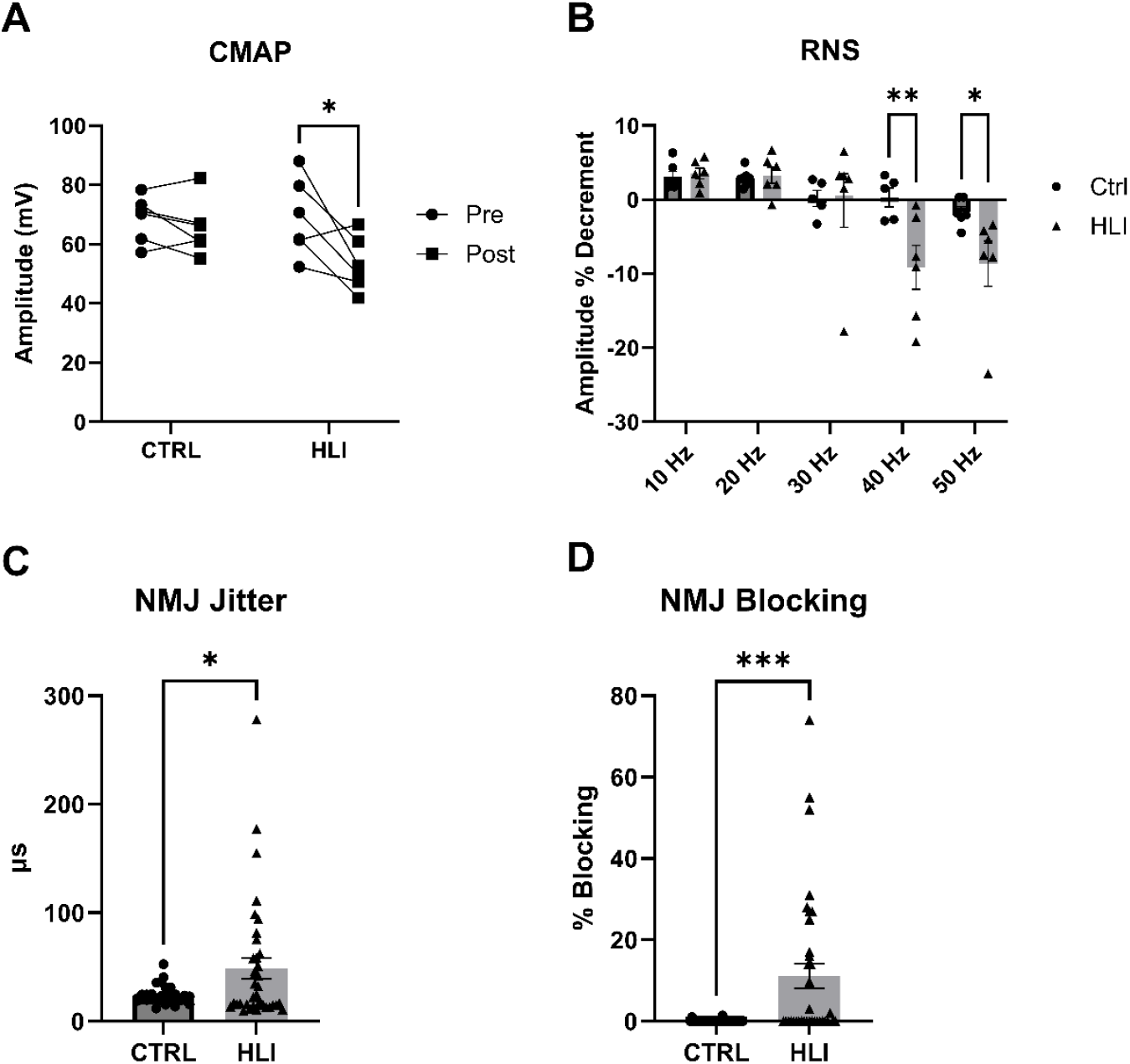
Deficits in neuromuscular excitability and NMJ transmission following 10 days HLI. A) Muscle excitability pre- and post-HLI. B) RNS decrement at 10, 20, 30,40 and 50Hz post-HLI. C) Quantification of jitter post-HLI, and D) quantification of blocking post-HLI. * p<0.05; ** p<0.01; *** p<0.001.

Next, SFEMG was employed to confirm our findings of NMJ transmission deficits. In HLI rats, we noted a marked 103% increase in NMJ transmission variability, termed jitter (Welch’s t test, 24 vs 48.74 µs, t=2.635, df=38.00, p=0.0121, **Fig. 4C**), and a striking presence of NMJ transmission failure, termed blocking (Welch’s t-test, 0.08095 vs 11.15 %, t=3.746, df=36.03, p=0.0006, **Fig. 4D**), relative to control rats.

## 4. Discussion

The present study demonstrates that just 10 days of unilateral hindlimb immobilization in young male rats induces neuromuscular changes consisting of muscle atrophy, reductions in muscle excitability and torque-generating capacity, and impaired NMJ transmission as evidenced by increased decrement on RNS and increased jitter and blocking on SFEMG. Taken together, these findings underscore that muscle weakness after short-term disuse cannot be explained by atrophy alone but also reflects robust impairments in neuromuscular excitability and NMJ transmission fidelity.

Ten days of unilateral HLI produced ∼20% muscle atrophy on average, consistent with prior reports showing that disuse induces rapid reductions in muscle mass within the first 1–2 weeks of immobilization ^4,5,20,21^. However, the decline in tetanic torque production, particularly when assessed by the torque-time integral, was modestly greater (∼23%) than the observed reduction in muscle size. This dissociation between muscle function and mass, while modest in our experiment utilizing young, healthy rats, has been demonstrated to be largely disproportional in aging^22-25^, suggesting that neural or synaptic mechanisms contribute substantially to disuse-induced weakness. Interestingly, despite the modest differences between muscle function and mass reported here, we still identify robust neuromuscular impairments as a hallmark feature of muscle disuse. In support, previous studies have implicated central drive, motor unit recruitment deficits, and NMJ morphological changes^7,10,26^, but direct evidence of NMJ transmission failure during short-term immobilization has been limited. Our electrophysiological results herein, in addition to our prior findings^14^, directly address this gap.

The marked increase in RNS decrement, NMJ jitter, and the occurrence of blocking observed with SFEMG provide direct electrophysiological evidence that short-term immobilization produces measurable transmission instability at the NMJ. These findings align with data from aging and neurodegenerative models, where NMJ instability has been documented as a key contributor to weakness out of proportion to muscle loss ^11,12,27-30^. Extending this concept to short-term disuse highlights the NMJ as an early and vulnerable site of dysfunction, even in otherwise healthy young muscle. Future work should expand these findings to the context of aging, where we hypothesize the disparity between loss of muscle function and mass following disuse becomes even more striking. It’s worth highlighting these findings are contradictory to some prior findings suggesting a lack of changes in NMJ morphology or transmission following disuse^31,32^. These discrepancies may be explained in the following ways: 1) method of muscle disuse used (i.e. immobilization versus hindlimb unloading), 2) muscle fiber-type composition of the muscle being studied (predominately type I versus type II fibers, which may be preferentially susceptible^33,34^), and 3) age of subjects. However, our findings are supported by a prior human study that also demonstrated the occurrence of NMJ jitter following unilateral cast immobilization in healthy young adults^35^. Mechanistically, inactivity has been demonstrated to result in morphological changes to the NMJ and may reduce synaptic activity-dependent signaling, leading to impaired acetylcholine release, receptor turnover, and structural destabilization of postsynaptic sites ^15,16^. Such changes would directly impair the fidelity of neuromuscular transmission and explain the functional deficits observed herein.

Importantly, these results provide strong translational implications. Patients undergoing muscle disuse whether due to sarcopenia, casting, prolonged bed rest, or critical illness, often exhibit profound weakness that exceeds the degree of measurable muscle atrophy^36-39^. Our findings provide mechanistic evidence that NMJ transmission failure contributes to this phenomenon, suggesting that therapeutic strategies aimed at preserving synaptic integrity and NMJ transmission could mitigate functional decline or enhance recovery following periods of disuse^29,30,40,41^.

This study is not without limitations. First, this work was conducted exclusively in young male rats, and the extent to which these findings generalize to females, across ages, or to humans, warrants further investigation. However, the findings reported herein recapitulate our previous work showing similar reductions in muscle excitability and pronounced RNS decrement in young female rats following ten days of HLI^14^, supporting NMJ transmission deficits as a feature of muscle disuse regardless of sex. Our electrophysiological assessments identify functional impairments in transmission, but a deeper understanding of the structural changes in the NMJ that underlie these deficits is still needed. Additionally, our study was restricted to a short-term disuse paradigm without recovery; the persistence or reversibility of NMJ dysfunction with reloading requires further investigation.

In conclusion, our results demonstrate that short-term immobilization induces not only muscle atrophy but also profound impairments in muscle excitability and neuromuscular transmission. By providing direct evidence of NMJ dysfunction following immobilization, this work shifts the understanding of disuse-related weakness from a largely myocentric process to one that includes NMJ transmission as a vital and underexplored contributing factor. These findings underscore the importance of targeting the NMJ, alongside muscle preservation, to mitigate the functional consequences of disuse.

